# A comparison of EEG encoding models using audiovisual stimuli and their unimodal counterparts

**DOI:** 10.1101/2023.11.16.567401

**Authors:** Maansi Desai, Alyssa M. Field, Liberty S. Hamilton

## Abstract

Communication in the real world is inherently multimodal. When having a conversation, typically sighted and hearing people use both auditory and visual cues to understand one another. For example, objects may make sounds as they move in space, or we may use the movement of a person’s mouth to better understand what they are saying in a noisy environment. Still, many neuroscience experiments rely on unimodal stimuli (visual only or auditory only) to understand encoding of sensory features in the brain. The extent to which visual information may influence encoding of auditory information and vice versa in natural environments is thus unclear. Here, we addressed this question by recording scalp electroencephalography (EEG) in 11 subjects as they listened to and watched movie trailers in audiovisual (AV), visual (V) only, and audio (A) only conditions. We then fit linear encoding models that described the relationship between the brain responses and the acoustic, phonetic, and visual information in the stimuli. We also compared whether auditory and visual feature tuning was the same when stimuli were presented in the original AV format versus when visual or auditory information was removed. We found that auditory feature tuning was similar in the AV and A-only conditions, and similarly, tuning for visual information was similar when stimuli were presented with the audio present (AV) and when the audio was removed (V only). In a cross prediction analysis, we investigated whether models trained on AV data predicted responses to A or V only test data as well as using the unimodal conditions for training. Overall, prediction performance using AV training and V test sets was similar to using V training and V test sets, suggesting that the auditory information has a relatively smaller effect on EEG. In contrast, prediction performance using AV training and A only test set was slightly worse than using matching A only training and test sets. This suggests the visual information has a stronger influence on EEG, though this makes no qualitative difference in the derived feature tuning. In effect, our results show that researchers may benefit from the richness of multimodal datasets, which can then be used to answer more than one research question.

## 1. Introduction

An important goal in auditory neuroscience is to understand the complex neural processes underlying speech perception, production, and language in humans. Traditional approaches to understanding brain processing of speech have focused on isolated phonemes, words, or various forms of evoked stimuli (e.g. clicks, chirps, pure tones) presented in controlled laboratory settings. With single word stimuli, researchers have shown that visual information can supplement auditory information in noisy settings, but these stimuli were tightly controlled to investigate the potential influence of lipreading [1]. In recent years, there has been growing interest in the use of naturalistic stimuli in speech research [2, 3, 4, 5]. Naturalistic stimuli refer to speech samples that are closer to real-world language use, such as continuous speech or conversational exchanges, rather than artificial, isolated words or sounds. This approach aims to capture the richness and complexity of human language and to better understand how speech is processed in our daily environment. Recently, studies have used continuous speech sentences [6, 7, 8, 9, 10] and audiobooks [11, 12, 13, 14, 15] to investigate speech encoding in the brain. These studies have found that neural activity measured through EEG is correlated with acoustic and linguistic stimuli, a phenomenon referred to as neural speech tracking. This approach can also be used to describe auditory feature selectivity, such as responses to specific phonological features like fricatives (e.g. /f/, /sh/, /s/, /v/, /z/, /zh/) or low back vowels (e.g /aa/), or specific responses to high frequency or low frequency spectral content in sounds [10, 9, 13, 8, 16]. However, many of these studies still use stimuli presented only through the auditory modality. Here, we wished to investigate how visual information influences auditory feature selectivity and vice versa when using naturalistic audiovisual stimuli and comparing them to their unimodal counterparts. Importantly, these stimuli were not designed to have a specific relationship between the auditory and visual features, but instead maintained their natural relationship.

Prior work using naturalistic stimuli have used video or audio recordings of natural scenes or events, such as social interactions, natural landscapes, or everyday activities [17, 9, 18, 19, 5, 20]. Studies have shown that our perception of audiovisual information is not simply the sum of individual sensory inputs but rather a dynamic process of integration, where multisensory information is combined and enhanced to improve the accuracy and speed of perceptual judgments [17, 21, 22]. However, it is unclear whether neural selectivity derived from a unimodal stimulus (audio or visual only) would be similar to selectivity derived from audiovisual stimuli in a less controlled experimental paradigm.

To address this question, we used EEG to record neural responses to speech and visual information in children’s movie trailers under three different conditions: audiovisual (AV), visual only (V), and audio only (A). Our first goal was to determine if auditory information was similarly represented when participants watched AV movies versus listening to the movie audio only (A). Then, we wanted to determine if neural responses to visual features were similarly represented in AV versus V conditions. Given our prior work showing that encoding models fit on audiovisual movie trailers could generalize to audio-only sentence stimuli [9], we hypothesized that the brain’s response to acoustic and phonetic features in AV would be similar to brain responses in A. This would mean that it would still be possible to use an audiovisual stimulus to investigate encoding of auditory features in the brain.

Using naturalistic audiovisual stimuli to generalize neural responses to more controlled listening paradigms may be fruitful for individuals who cannot tolerate lengthy and tedious tasks [18, 9]. Because of the low signal-to-noise ratio of EEG, long experimental sessions are often required when investigating brain processing of speech. For children and clinical populations, long experimental sessions may not be tolerable at all if the tasks are also considered to be boring or unpleasant. Thus, our research is partially motivated by attempts to create tasks that will maximize data quality and amount of data needed to collect for an experimental paradigm [18]. A recent study found that tracking of acoustic and linguistic speech features was feasible using both continuous speech sentences from both the Texas Instruments Massachusetts Institute of Technology (TIMIT) speech corpus [23] and movie trailers [9, 18]. This study also showed that regressing out the visual information did not impact the ability to predict neural responses to speech.

The purpose of the current study is to understand how information in one sensory modality (auditory or visual) is encoded by the brain during audiovisual movie watching versus unimodal presentation of these stimuli during EEG recordings. The results from this study will provide practical information for interpreting the results of experiments using naturalistic multimodal stimuli with the overarching goal of utilizing stimuli that are jointly acoustically and visually rich, while allowing experimenters to interpret multimodal or unimodal information depending on the research question at hand.

## 2. Methods

### 2.1. Data

#### 2.1.1. Participants

11 native-English participants with normal hearing (5M, 5F, 1 N/B, age 18-31, mean age: 25.6 *±* 3.95 years) were recruited to participate. The experimental procedures were approved by The University of Texas at Austin Institutional Review Board. All participants gave written informed consent to participate in the task. Pure tone thresholds (250-8000 Hz) and speech-in-noise hearing tests (QuickSIN, Interacoustics) were administered to ensure that the participants had normal hearing (*<* 25 dB HL thresholds for pure tone and *<* 3 dB SNR loss for QuickSIN). All participants had normal or corrected-to-normal vision.

#### 2.1.2. Experimental Design

EEG participants listened to and watched children’s movie trailers for approximately 1 hour, taken from a previously published dataset (https://osf.io/p7qy8/) [9]. These stimuli contain overlapping speech, music, background sounds, and visual information. The movie trailer stimuli were hand transcribed to reflect the onset and offset of phoneme and word-level boundaries using ELAN (https://archive.mpi.nl/tla/elan) and a modified version of the Penn Phonetics forced aligner, FAVE align (https://zenodo.org/record/9846). Timings were manually corrected using Praat by four raters to ensure reliability. In addition to the linguistic information, the onset and offset of scene changes was marked.

During the task, all EEG participants listened to and watched eight movie trailers in an audiovisual condition (AV: listening and watching), a visual only condition (V: only watching with no audio), and a listening only condition (A: only listening to the audio and staring at a fixation cross). AV, V, and A trials were presented in pseudo-random order. The task was presented through an iPad running custom software written in Swift (version 5.3.2; https://developer.apple.com/swift/). The iPad was placed on a table at precisely 23 inches from the participant, measured from the nasion to the top of the iPad. Videos were rendered at 1280 *×* 720-pixel movies at 24 frames/s. The audio was sampled at 44.1 kHz. Each movie trailer for AV, V, and A conditions was presented once.

#### 2.1.3. Data Acquisition

High-density EEG data were collected using a standard 64-channel montage (EasyCap) at a sampling rate of 25 kHz using the BrainVision actiCHamp system (Brain Products). Impedance for all channels was kept below 15 *k*Ω. An additional two channels were used to measure vertical and horizontal electrooculography (EOG) to remove ocular artifacts during preprocessing. The audio from the task was synchronized with the neural data using a StimTrak stimulus processor (Brain Products). The visual only condition contained a short notification sound which was used to synchronize the onset of the stimulus with the EEG data.

#### 2.1.4. EEG Preprocessing

EEG and EOG electrode timecourses were downsampled to 128 Hz using the BrainVision Analyzer 2.1 software. Subsequent preprocessing steps were conducted offline using MNE-Python [24]. EEG channels were re-referenced to the average of the mastoid channels (TP9 and TP10) and then notch filtered at 60 Hz to remove line noise. The data were bandpass filtered between 1-15 Hz using a zero-phase, noncausal bandpass FIR (finite impulse response) filter (Hamming window, 0.0194; passband ripple with 53 dB stopband attenuation, −6 dB falloff). Manual artifact rejection was conducted to remove large non-biological artifactual movement, and no more than 10% of the data were removed. Independent component analysis (ICA) with 64 specified components was used to remove blinks and saccades. Finally, EEG data were epoched to the onset and offset of the acoustic stimuli (for AV and A conditions) and the short notification sounds (V only). A customized Python script used a match filter procedure [25] to convolve the stimulus waveform with the recorded audio from the EEG experiment. Once epoched, the onset of each trailer for each condition (AV, A, V) was used to identify the acoustic and linguistic onset information from the Praat textgrids [26].

#### 2.1.5. Auditory and Visual Feature Extraction

For auditory features, we extracted the acoustic envelope, pitch, and phonological features from our stimuli. For phonological features, we created a binary matrix to indicate the timing of all phonemes based on place and manner of articulation for each movie trailer (e.g. sonorant, obstruent, fricative, etc.) [10, 9]. The matrix contained a 1 at the onset of the phonological feature and 0 for all other times. The acoustic envelope of the stimuli was extracted using the Hilbert-transform of the waveform followed by a low-pass filter (third-order Butterworth filter; cutoff frequency, 25 Hz) and downsampling to 128 Hz. Lastly, the absolute pitch was calculated using an autocorrelation method implemented in Parselmouth [27], a python package that inter-faces with Praat [26], with a 7.8 ms window (1/128 Hz) and 50-300 Hz F0 range.

While the movie trailers are acoustically rich, these stimuli also contain visual information. To investigate the influence of these visual features on audiovisual speech encoding, we fit encoding models using scene cuts and Gabor wavelet filters in addition to the acoustic features above. Scene cut information was taken from Praat textgrids, which contained hand-annotated onsets of scene changes. Gabor wavelet filters were calculated using a nonlinear Gabor motion energy filter bank [28] to capture varying visual motion for every frame of each movie trailer. A total of 2139 filters were calculated and a principal component analysis decomposed these features into 10 components to reduce the dimensionality of the feature matrix.

#### 2.1.6. Linear Encoding Models

Encoding models (known as temporal response functions (TRFs), multivariate temporal receptive fields (mTRFs), or linear regression) are a method for describing the statistical relationship between a stimulus (e.g. visual or auditory feature representations) and neural response [29, 8, 9, 30, 10, 31, 13]. We fit mTRF models to all 64 channels from all EEG participants, using separate auditory or visual feature models (Equation 1).

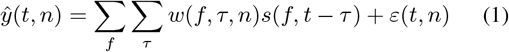

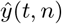 (*t, n*) represents the neural response for each EEG channel, *n*, at time, *t. w*(*f, τ, n*) is a matrix of weights fit from the model for a given feature representation (*f*), at each channel (*n*), and time delay (*τ*). Weights were fit using ridge regression on a subset of training data (Equation 2).

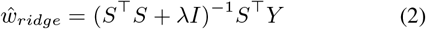

The regularization parameter (*λ*) was estimated using a separate cross-validation procedure, where we tested 15 values of *λ* between 10^2^ and 10^8^. This was performed for 100 bootstrap iterations of portions of the training set. The time delay (*τ*) for each model fit was between 0 and 600 ms. *s*(*f, t* − *τ*) is the stimulus input used to fit the encoding model for a given feature representation for the specified time delay. *ε*(*t, n*) are the residual errors for the linear regression. The test set data consisted of the average of the same two unique movie trailers for each respective condition (AV, V, A). The remaining 6 trailers were used as training data. For each encoding model using separate auditory or visual features, the performance was evaluated by calculating the correlation (*r*) between the predicted EEG and the actual held out EEG.

#### 2.1.7. Multimodal and unimodal model comparisons

We compared the performance of models trained on multimodal audiovisual data to the performance of models trained on unimodal (auditory only or visual only) stimulus presentations. In addition, we compared the structure of the weights and derived feature tuning across EEG channels when using these different stimuli. Topographical maps were plotted to show the spatial distribution of encoding model performance for each condition type.

#### 2.1.8. Cross Prediction Analysis

Part of our goal in this study was to determine whether stimulus selectivity was similar in unimodal (auditory or visual only) conditions compared to audiovisual conditions. To determine whether receptive field models generalized across condition type, we conducted a cross-prediction analysis to predict responses to auditory information based on the audiovisual stimulus-derived receptive fields and compared these to predictions to those derived from auditory only stimulus-derived receptive fields. We also performed the same analysis to compare predictions of neural responses to visual information when using the audiovisual data versus the visual only data. If the audiovisual stimulus does not significantly affect the structure of receptive fields, we should see prediction performance that is similar for unimodal versus multimodal conditions.

When predicting auditory responses from audiovisual weights, we used all speech features (phonological features, the acoustic envelope, and pitch). When predicting visual responses from audiovisual weights, we used all visual features (Gabor wavelet filters and scene cuts). We then used the weights calculated from the audiovisual condition as the training set to predict the neural responses of the respective speech (auditory condition) or visual (visual condition) features in the movie trailer EEG data. That is, using pre-trained models from the audiovisual condition, we predicted responses to the auditory or visual only condition as the test set. We then compared the model performance for the cross-condition analysis to the model performance for the original within-condition analysis, where training and test data came from the same condition type.

### 2.2. Data and Code Availability

Code available at https://github.com/HamiltonLabUT/EEG-encoding-audiovisual.

Data available at https://osf.io/tgbk7/.

## 3. Results

### 3.1. Auditory feature representations in the brain are similar for uni- and multimodal stimuli

Encoding models provide a statistical metric to map the relationship between the brain’s response and continuous stimuli. Here, we test the encoding of audio and visual information as represented in these EEG signals.

We trained on 64-channel EEG data responses using a combination of 14 phonological features, the acoustic envelope, and pitch. These features have been used in previous work to investigate acoustic and linguistic selectivity to continuous speech stimuli [13, 8, 10, 9]. Model performance was assessed by comparing the linear regression correlation value (*r*) between the actual and predicted EEG for each condition (A versus AV). We hypothesized that encoding of auditory features between A and AV would generate comparable model performance. Model performance was similar across all subjects for both A (*r*_*max*_ = 0.13, *r*_*avg*_ = 0.02) and AV (*r*_*max*_ = 0.22, *r*_*avg*_ = 0.02) conditions (Figure 1A). Individual model performance for each EEG channel in all subjects was compared for A vs. AV, with most points falling along the unity line. The structure of the weights for AV versus A looked similar, with broad selectivity and enhancement of voltage changes to linguistic and acoustic features for both conditions between 0.0 and 0.25 seconds post stimulus onset. Topographical maps show that the highest model performance was primarily concentrated in frontotemporal regions of the scalp for both AV and A models (Figure 1B). Post-hoc Wilcoxon signed-rank tests showed the AV condition correlations were significantly smaller than the A condition (*W* = 113446.0, *p* = 0.0488). The effect size was extremely small (*median*_*AV −A*_ = − 0.001), however, so this difference is unlikely to have any practical effect.

**Figure 1:**
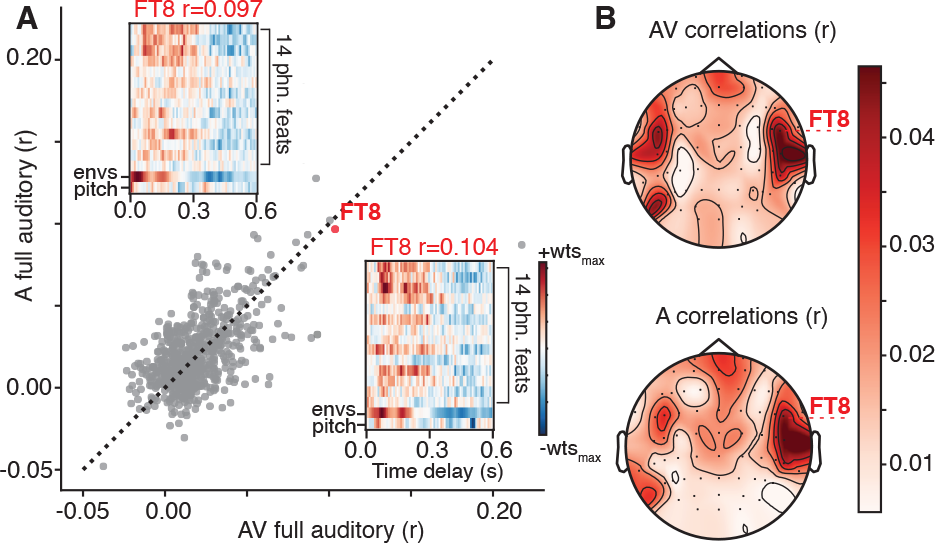
The auditory feature model used a combination of phonological features, acoustic envelope, and pitch to predict EEG responses to auditory information in AV and A conditions. A) Scatter plot shows the model comparison (r-value) between AV and A conditions. Each gray dot represents the encoding model performance in a single channel in a single participant. Channel FT8 is shown for subject MT0033 with the corresponding weight matrices for each condition (A condition: top left, AV condition: bottom right) and associated correlation value for the channel. B) Average correlation values for AV and A condition plotted on topographical map and averaged across all participants (N=11). Topography of selective channels was similar for AV and A.

This first analysis demonstrated comparable model performance for speech encoding when using multiple acoustic and linguistic features for the AV versus A only condition. While the results suggested that similar auditory feature tuning can be derived from A only and AV stimuli, we also wished to investigate the converse – whether similar visual feature tuning could be derived from V only and AV stimuli.

### 3.2. Visual feature representations in the brain are not impacted by the presence of auditory information

Our previous analysis showed that auditory feature encoding was comparable between AV and A only conditions. However, the visual features themselves are a prominent perceptual experience when engaging with movie trailers. So how much does auditory information impact visual encoding, if at all? Previous work has demonstrated that the addition of visual information contributed to a significant portion of variance explained, but does not impact the ability to encode speech information [9]. To better understand the converse relationship, we trained models for all 64 EEG channels across all participants using a combination of Gabor wavelet features and a binary feature vector of scene cut information [28, 9]. The Gabor filters were reduced to 10 principal components to mitigate computational complexity when fitting the encoding models, as described in [9]. In addition, we included the timing of scene cuts in this linear regression model.

Similar to the auditory feature encoding, we hypothesized that visual information should be similarly represented in the visual (V) only condition despite the presence of auditory information from the audiovisual condition. We compared correlation (*r*) values between actual and predicted EEG data from V and AV across all participants using 11 visual features (gabor wavelets and scene cuts). Encoding model performance was similar for the V (*r*_*max*_ = 0.28, *r*_*avg*_ = 0.08) and AV (*r*_*max*_ = 0.26, *r*_*avg*_ = 0.08) conditions (Figure 2A). Individual channel encoding models were distributed across the unity line. The structure of the weights was visually similar, as shown for an example single channel (P4) in one participant (MT0029). In this example, the scene cuts induce similar voltage changes across time delays in both V and AV conditions. Model performance for AV (*r* = 0.222) and V (*r* = 0.223) conditions were also highly similar for this channel. Topographical maps of the average correlation values from all subjects across all 64 EEG channels show that the highest *r*-values for visual encoding models were concentrated in occipital regions of the scalp (Figure 2B), regardless of AV or V presentation. Post-hoc Wilcoxon signed-rank tests showed that the AV condition correlations were statistically significantly smaller than the V condition (*W* = 113362.0, *p* = 0.0471). Similar to the AV and A comparison, the effect size was again minuscule (*median*_*AV −V*_ = − 0.00235).

**Figure 2:**
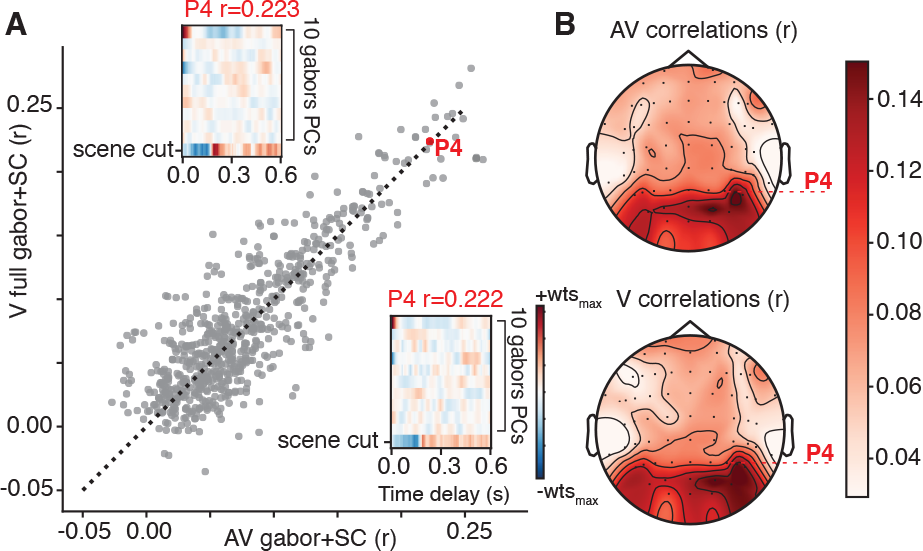
Visual feature model using a combination of 10 Gabor wavelet filter principal components (PCs) and scene cut (SC) information. A) Scatter plot shows the model comparison (r-value) between AV and V conditions. Each gray dot represents the encoding model performance in a single channel in a single participant. An example channel, P4, is shown for subject MT0029 with the corresponding weight matrices for each condition and associated model performance (correlation value) for the channel. B) Grand average model performance (correlation values) for AV and V condition plotted on a topographical map show similar spatial distribution of good model performance regardless of condition.

In addition to using visual features to compare neural responses between the AV and V conditions, we fit an encoding model just using the 14 phonological features and compared model performance correlation values (*r*) between the AV and V conditions. The purpose of this analysis was to see whether auditory or phonological information might be inferred from the visual information, even when it was absent in the V only condition. In this case, although sound was technically absent in the V condition, we used the timings of the phonological features from the AV condition, which would be the same. We found that models predicting responses to the inferred phonological features in the V condition performed below chance (*r*_*max*_ = 0.088, *r*_*avg*_ = − 0.0015) compared to models predicting responses to this linguistic information the AV condition (*r*_*max*_ = 0.11, *r*_*avg*_ = 0.017) and A condition (*r*_*max*_ = 0.13, *r*_*avg*_ = 0.019). These results demonstrate that phonological features are not obviously represented in the V condition compared to AV or A. This difference may be due to the relative independence of the auditory and visual information in this stimulus set, unlike studies assessing lip-reading where most audiovisual information is congruent [32, 33, 34, 35]. While previous results compared auditory and visual feature encoding in AV, V, A conditions, the impact of combining specific visual and auditory information, such as congruent and incongruent mouth movement with phonological feature encoding in AV and V conditions is still unclear.

### 3.3. Cross prediction analysis

In the previous analysis, we showed that prediction performance was comparable for unimodal and multimodal encoding models (Fig. 1 and 2). In addition, the receptive field models derived from data in both unimodal and multimodal conditions exhibited similar selectivity. Still, this comparison does not address whether models fit on one data type (uni- or multimodal) would predict responses to the other type of dataset.

To address this, we conducted further analyses to determine whether encoding models trained using audiovisual could generalize and accurately predict neural responses in the unimodal conditions (A only or V only). The motivation behind this analysis was to test whether the presence of visual information significantly influenced predictions for the auditory modality compared to when information is presented in isolation. This would allow us to determine if auditory receptive fields or visual receptive fields derived from audiovisual information are a reliable estimate of how the brain responds in the unimodal context. This would allow researchers to consider using more naturalistic multimodal stimuli in their experiments, even if their research questions related to unimodal sensory encoding.

For the cross-prediction analysis, we manipulated the training set type (uni vs. multimodal) and measured prediction performance for data from each unimodal test set. For example, we used the acoustic model weights (phonological features, pitch, and acoustic envelope) from the A-only condition to predict responses to test set data from A-only and test set data from AV neural data. Similarly, we used the combination of Gabor wavelet filters and scene cuts from the V-only condition to predict responses to untrained V-only and AV EEG data. For both of these cross-predictions, we compared the performance of the model trained on the unimodal condition and tested on the same unimodal condition (e.g. predicting unimodal A or V from unimodal A or V training set data) or trained on the multimodal condition (e.g. predicting A or V from AV training set data).

Results of this cross-prediction comparison are shown in Figure 3. Correlation values below the unity line indicate electrodes for which the within-condition model performance was better, and those above the unity line indicate better cross-condition performance. A regression line was plotted for both stimulus comparisons to determine how correlated the individual encoding model was with the cross-predicted model performance. Overall, using the same condition type for training and testing (e.g., predict A test data from A training data or predict V test data from V training data) resulted in better model performance than cross-predictions (e.g. predict A test data from AV training data, or predict V test data from AV training data). However, the response to one condition (A or V) could be modeled from the AV condition and was significantly correlated (r=0.41, p¡0.001 for the auditory comparisons in Fig 3A, and r=0.902, p¡0.001 for the visual comparisons in Fig. 3B). Similar to the previous results, predicting auditory responses from A-only and AV data resulted in lower correlation values (Figure 3A) compared to the predicting visual responses from V-only and AV data (Figure 3B). In previous work, we found that the model performance for a visual feature model was greater than models using acoustic information when using the movie trailer in its original multimodal form. [9]. The results here in Figure 3A and 3B also corroborate such findings for unimodal stimuli. Still, we found that even when using the multimodal (AV) condition data, it was still possible to encode neural responses to A or V-only stimulus information. This suggests that audiovisual stimuli provide generalizable results to unimodal stimuli, with the added benefit of providing a rich source of information for asking multiple research questions. We hypothesized that the correlation from the cross-prediction analysis may arise from highly salient scene cut information, which could contribute to a larger dynamic visual response. To test if the model performance from the visual information differed based on feature representation, we conducted a cross prediction analysis using the gabor only or scene cut only model from the V only condition and tested on the same visual feature in the AV condition (Figure 3C). We found that the scene cut information was concentrated along the unity line, suggesting that the scene cut information was equally modeled in the V-only and AV conditions. The gabor only information has slightly better model performance in the V-only condition. Finally, we correlated each channel for each participant between AV and A and AV and V conditions and found that more of the occipital channels were correlated for the visual conditions. However, the frontal and temporal regions across the scalp were not correlated for the AV and A condition (Figure 3D). This suggests that the visual information has an outsized effect on the EEG itself in both visual and audiovisual conditions, although the weights are correlated between unimodal and multimodal conditions (Figure 1-2).

**Figure 3:**
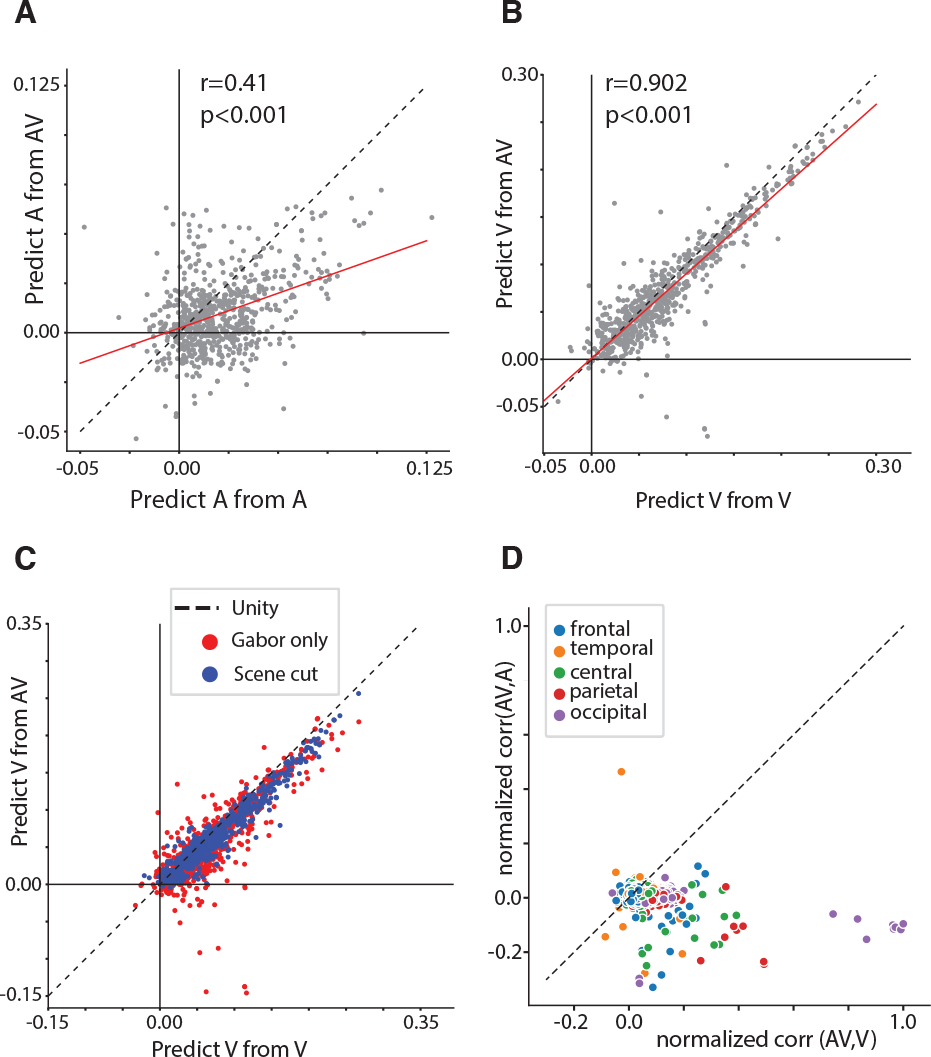
Cross prediction analysis shows that responses are generalizable between unimodal and multimodal stimulus information, with stronger generalizability for visual information compared to auditory. A) Model performance for audio-only (A) test sets with A-only training data (x-axis) or audiovisual (AV) training data (y-axis), calculated as the linear correlation between predicted and actual held out EEG test data. Each dot represents an individual electrode for an individual EEG subject (64 channels x 11 participants). Dashed black line = unity line; red line = regression line. B) Model performance for visual-only (V) test set with V-only training data or AV training data. Similar model performance was observed for both within- and cross-condition predictions, though this relationship was stronger between V and AV. C) Model performance for comparing visual only responses using either scene cuts or only gabor feature representations with the individual feature in the audio-visual condition. D) Normalized correlation coefficient between each EEG channel for audiovisual and visual only conditions and audiovisual and audio only conditions. Overall, single trial bandpass filtered EEG (input to the model) was more correlated between the AV and V only conditions as compared to the AV and A only conditions, suggesting a strong influence of visual information on the EEG signals.

## 4. Discussion

The use of naturalistic audiovisual stimuli in EEG research has become increasingly important in understanding the complex relationship between sensory modalities, while simultaneously providing a unique and valuable tool for investigating neural processing in a more ecologically valid context. Using stimuli that are more engaging to investigate neural responses to speech or visual information improves the participant’s experience with the experimental paradigm, leading them to feel less fatigued and discontent [2]. Here, we designed an experiment to identify if auditory and visual information were differently encoded in natural stimuli in three varying conditions: AV, V, A. We show that AV training data can be used to derive both auditory and visual receptive fields that are comparable to those achieved using unimodal stimuli. In other words, for these movie stimuli, it is possible to identify neural responses to speech using the AV condition without worrying about the visual feature interfering with the linear regression model performance or receptive field weights. Our cross-prediction analysis demonstrated that we could independently uncover unimodal feature tuning even when using a naturalistic audiovisual stimulus set. While we found similar receptive field tuning for some electrodes when comparing A-only and AV and comparing V-only with AV, predicting neural responses to visual information was similarly robust when using AV training data versus V training data. In contrast, predicting neural responses to auditory information from the AV condition was possible but not as robust as the model performance for AV versus V data. The results from this study and previous work using these stimuli [9] showed that while it is possible to use an acoustically rich audiovisual stimuli such as this one to encode neural responses to speech, the visual information does contain a large presence in the EEG signal. As such, the results from the cross-prediction analysis in Figure 3 may demonstrate that the EEG signal captures more visual information compared to the auditory information when using multimodal naturalistic stimuli. Ultimately, the relatively larger influence of visual information on the models as compared to auditory information suggests further studies are needed to disentangle these relationships. We provide motivation for replacing more controlled naturalistic speech-only experimental paradigms with tasks that include multisensory information and are better representative of speech encoding in our daily environment. Additionally, using AV stimuli allows for addressing multiple research questions with the same rich dataset.

## 5. Conclusions

We fit encoding models to predict EEG responses to auditory features (phonological features, the acoustic envelope, pitch) and visual features (gabor wavelet and scene cut) using AV, A-only, and V-only conditions. We found that presenting an audiovisual stimulus resulted in similar encoding model weights to those derived from auditory or visual only stimuli. However, researchers should be aware of strong influences of visual evoked potentials on the ongoing EEG. Overall, movie trailers can be used as naturalistic audiovisual stimuli to better understand neural encoding of speech or visual information.

## 6. Acknowledgements

The authors would like to thank all of the EEG subjects. This work was supported by a grant from the Texas Speech-Language Hearing Foundation. We would also like to thank Alexander Huth for helpful comments on this manuscript.

## References

[1] M. Ozker, I. M. Schepers, J. F. Magnotti, D. Yoshor, and M. S. Beauchamp, “A double dissociation between anterior and posterior superior temporal gyrus for processing audiovisual speech demonstrated by electrocorticography,” Journal of cognitive neuroscience, vol. 29, no. 6, pp. 1044–1060, 2017.

[2] L. S. Hamilton and A. G. Huth, “The revolution will not be controlled: natural stimuli in speech neuroscience,” Language, cognition and neuroscience, vol. 35, no. 5, pp. 573–582, 2020.

[3] S. Sonkusare, M. Breakspear, and C. Guo, “Naturalistic stimuli in neuroscience: critically acclaimed,” Trends in cognitive sciences, vol. 23, no. 8, pp. 699–714, 2019.

[4] Y. Zhang, J.-H. Kim, D. Brang, and Z. Liu, “Naturalistic stimuli: A paradigm for multiscale functional characterization of the human brain,” Current opinion in biomedical engineering, vol. 19, p. 100298, 2021.

[5] P. J. Matusz, S. Dikker, A. G. Huth, and C. Perrodin, “Are we ready for real-world neuroscience?” pp. 327–338, 2019.

[6] E. F. Chang, J. W. Rieger, K. Johnson, M. S. Berger, N. M. Barbaro, and R. T. Knight, “Categorical speech representation in human superior temporal gyrus,” Nature neuroscience, vol. 13, no. 11, pp. 1428–1432, 2010.

[7] C. Tang, L. Hamilton, and E. Chang, “Intonational speech prosody encoding in the human auditory cortex,” Science, vol. 357, no. 6353, pp. 797–801, 2017.

[8] L. S. Hamilton, E. Edwards, and E. F. Chang, “A spatial map of onset and sustained responses to speech in the human superior temporal gyrus,” Current Biology, vol. 28, no. 12, pp. 1860–1871, 2018.

[9] M. Desai, J. Holder, C. Villarreal, N. Clark, B. Hoang, and L. S. Hamilton, “Generalizable eeg encoding models with naturalistic audiovisual stimuli,” Journal of Neuroscience, vol. 41, no. 43, pp. 8946–8962, 2021.

[10] N. Mesgarani, C. Cheung, K. Johnson, and E. F. Chang, “Phonetic feature encoding in human superior temporal gyrus,” Science, vol. 343, no. 6174, pp. 1006–1010, 2014.

[11] H. Akbari, B. Khalighinejad, J. L. Herrero, A. D. Mehta, and N. Mesgarani, “Towards reconstructing intelligible speech from the human auditory cortex,” Scientific reports, vol. 9, no. 1, p. 874, 2019.

[12] M. P. Broderick, A. J. Anderson, G. M. Di Liberto, M. J. Crosse, and E. C. Lalor, “Electrophysiological correlates of semantic dissimilarity reflect the comprehension of natural, narrative speech,” Current Biology, vol. 28, no. 5, pp. 803–809, 2018.

[13] G. M. Di Liberto, J. A. O’sullivan, and E. C. Lalor, “Low-frequency cortical entrainment to speech reflects phoneme-level processing,” Current Biology, vol. 25, no. 19, pp. 2457–2465, 2015.

[14] L. Gwilliams, J.-R. King, A. Marantz, and D. Poeppel, “Neural dynamics of phoneme sequences reveal position-invariant code for content and order,” Nature communications, vol. 13, no. 1, p. 6606, 2022.

[15] M. Gillis, J. Vanthornhout, J. Z. Simon, T. Francart, and C. Brodbeck, “Neural markers of speech comprehension: measuring eeg tracking of linguistic speech representations, controlling the speech acoustics,” Journal of Neuroscience, vol. 41, no. 50, pp. 10 316–10 329, 2021.

[16] L. S. Hamilton, Y. Oganian, J. Hall, and E. F. Chang, “Parallel and distributed encoding of speech across human auditory cortex,” Cell, vol. 184, no. 18, pp. 4626–4639, 2021.

[17] M. J. Crosse, G. M. Di Liberto, and E. C. Lalor, “Eye can hear clearly now: inverse effectiveness in natural audiovisual speech processing relies on long-term crossmodal temporal integration,” Journal of Neuroscience, vol. 36, no. 38, pp. 9888–9895, 2016.

[18] M. Desai, A. M. Field, and L. S. Hamilton, “Dataset size considerations for robust acoustic and phonetic speech encoding models in eeg,” Frontiers in Human Neuroscience, vol. 16, 2022.

[19] R. Thézé, M. A. Gadiri, L. Albert, A. Provost, A.-L. Giraud, and P. Mégevand, “Animated virtual characters to explore audio-visual speech in controlled and naturalistic environments,” Scientific reports, vol. 10, no. 1, p. 15540, 2020.

[20] J. Berezutskaya, M. J. Vansteensel, E. J. Aarnoutse, Z. V. Freudenburg, G. Piantoni, M. P. Branco, and N. F. Ramsey, “Open multimodal ieeg-fmri dataset from naturalistic stimulation with a short audiovisual film,” Scientific Data, vol. 9, no. 1, p. 91, 2022.

[21] R. A. Stevenson, M. T. Wallace, and N. Altieri, “The interaction between stimulus factors and cognitive factors during multisensory integration of audiovisual speech,” Frontiers in psychology, vol. 5, p. 352, 2014.

[22] S. Ten Oever, A. T. Sack, K. L. Wheat, N. Bien, and N. Van Atteveldt, “Audio-visual onset differences are used to determine syllable identity for ambiguous audio-visual stimulus pairs,” Frontiers in psychology, vol. 4, p. 331, 2013.

[23] J. S. Garofolo, L. F. Lamel, W. M. Fisher, J. G. Fiscus, and D. S. Pallett, “Darpa timit acoustic-phonetic continous speech corpus cd-rom. nist speech disc 1-1.1,” NASA STI/Recon technical report n, vol. 93, p. 27403, 1993.

[24] A. Gramfort and L. M. et al., “Meg and eeg data analysis with mne-python,” Frontiers in Neuroscience, vol. 7, p. 267, 2013.

[25] G. Turin, “An introduction to matched filters,” IRE transactions on Information theory, vol. 6, no. 3, pp. 311–329, 1960.

[26] P. Boersma and D. Weenink, “Praat: doing phonetics by computer [Computer program],” Version 6.1.38, retrieved 2 January 2021 http://www.praat.org/, 2021.

[27] Y. Jadoul, B. Thompson, and B. de Boer, “Introducing Parselmouth: A Python interface to Praat,” Journal of Phonetics, vol. 71, pp. 1–15, 2018.

[28] S. Nishimoto, A. T. Vu, T. Naselaris, Y. Benjamini, B. Yu, and J. L. Gallant, “Reconstructing visual experiences from brain activity evoked by natural movies,” Current biology, vol. 21, no. 19, pp. 1641–1646, 2011.

[29] M. J. Crosse, G. M. Di Liberto, A. Bednar, and E. C. Lalor, “The multivariate temporal response function (mtrf) toolbox: a matlab toolbox for relating neural signals to continuous stimuli,” Frontiers in human neuroscience, vol. 10, p. 604, 2016.

[30] C. R. Holdgraf, J. W. Rieger, C. Micheli, S. Martin, R. T. Knight, and F. E. Theunissen, “Encoding and decoding models in cognitive electrophysiology,” Frontiers in systems neuroscience, vol. 11, p. 61, 2017.

[31] F. E. Theunissen, K. Sen, and A. J. Doupe, “Spectral-temporal receptive fields of nonlinear auditory neurons obtained using natural sounds,” Journal of Neuroscience, vol. 20, no. 6, pp. 2315–2331, 2000.

[32] A. Begau, L.-I. Klatt, E. Wascher, D. Schneider, and S. Getzmann, “Do congruent lip movements facilitate speech processing in a dynamic audiovisual multi-talker scenario? an erp study with older and younger adults,” Behavioural Brain Research, vol. 412, p. 113436, 2021.

[33] M. Bourguignon, M. Baart, E. C. Kapnoula, and N. Molinaro, “Lip-reading enables the brain to synthesize auditory features of unknown silent speech,” Journal of Neuroscience, vol. 40, no. 5, pp. 1053–1065, 2020.

[34] M. J. Crosse, J. S. Butler, and E. C. Lalor, “Congruent visual speech enhances cortical entrainment to continuous auditory speech in noise-free conditions,” Journal of Neuroscience, vol. 35, no. 42, pp. 14 195–14 204, 2015.

[35] F. Bröhl, A. Keitel, and C. Kayser, “Meg activity in visual and auditory cortices represents acoustic speech-related information during silent lip reading,” eneuro, vol. 9, no. 3, 2022.

